# Two forms of isometric muscle function: Interpersonal motor task supports a distinction between a holding and a pushing isometric muscle action

**DOI:** 10.1101/2020.08.17.253633

**Authors:** Laura V Schaefer, Frank N Bittmann

**Affiliations:** Section Regulative Physiology and Prevention, Department Sports and Health Sciences, University of Potsdam, Potsdam, Germany

**Keywords:** Two forms of isometric muscle action, holding isometric muscle action (HIMA), pushing isometric muscle action (PIMA), interpersonal motor task, mechanomyography, mechanotendography

## Abstract

In sports and movement sciences isometric muscle function is measured by pushing against a stable resistance. However, subjectively one can hold or push isometrically. Several investigations suggest a distinction of those forms. The aim of this study was to investigate, whether or not these two forms of isometric muscle action can be distinguished by objective parameters in an interpersonal setting. 20 subjects were grouped in 10 same sex pairs, in which one partner should perform the pushing isometric muscle action (PIMA) and the other partner executed the holding isometric muscle action (HIMA). The partners were coupled by an interface including a strain gauge and an acceleration sensor. The mechanical oscillations of the triceps brachii (MMGtri) muscle, its tendon (MTGtri) and the abdominal muscle (MMGobl) were recorded by piezoelectric-sensor-based measurement system (mechanomyography (MMG); mechanotendography (MTG)). Each partner performed three 15s (80% MVIC) and two fatiguing trials (90% MVIC) during PIMA and HIMA, respectively (tasks changed in the couple). Regarded parameters to compare PIMA and HIMA were (1) the mean frequency, (2) the normalized mean amplitude, (3) the amplitude variation, (4) the power in the frequency range of 8 to 15 Hz and (5) a special power-frequency ratio and the number of task failures during HIMA or PIMA (partner who quit the task).

A “HIMA failure” occurred in 87.5% of trials (*p*<0.000). No significant differences between PIMA and HIMA were found for the mean frequency and normalized amplitude. The MMGobl showed a significantly higher values of the amplitude variation (15s: *p* 0.013; fatigue: *p*=0.007) and of the power-frequency-ratio (15s: *p* = 0.040; fatigue: *p* = 0.002) during HIMA and a higher power in the range of 8 to 15 Hz during PIMA (15s: *p*=0.001; fatigue: *p*=0.011). MMGtri and MTGtri showed no significant differences.

Based on the findings it is suggested that a holding and a pushing isometric muscle action can be distinguished objectively, whereby a more complex neural control is assumed for HIMA.

## Introduction

Usually isometric muscle action is measured by pushing against a stable resistance. Subjectively, however, one can perform isometric muscle action by pushing against a stable resistance or by resisting an object, thus, in a holding manner. For example, during arm wrestling, one can pursue the strategy to push against the partner until he or she gives in or one can try to hold as long as possible until the partner cannot push anymore. In both cases, no motion occurs as long as both partners are able to maintain the reaction force, thus, the muscle length and joint angle stay stable. In a recent study, we suggested that those forms are referred to as the pushing isometric muscle action (PIMA) and the holding isometric muscle action (HIMA), respectively [1]. Until now, the holding form is rarely considered in movement or sport sciences. Garner et al. hypothesized that two forms of isometric muscle action exist [2]. They rejected the hypothesis, since no differences in the amplitude or frequency of Electromyography (EMG) of the soleus and gastrocnemius muscles appeared during the performance of HIMA and PIMA. Furthermore, the research group around Enoka [3–7] investigated those assumed different forms of isometric muscle action and denoted them position task (HIMA) or force task (PIMA). Thereby, the position task (HIMA) was performed by holding an inertial load and maintaining a constant position of the limb, which was freely movable. During the force task (PIMA) a force level (same intensity as during the position task) should be maintained by pushing against a stable resistance. The main finding was that the time to task failure was significantly longer during the force task compared to the position task for the elbow flexor muscles at intensities of 15% of the maximal voluntary isometric contraction (MVIC) [4,5,7]. However, those results were only found at low intensities of 15% to 30% of the MVIC and with the forearm positioned horizontally. With higher force intensities of 45% and 60% of the MVIC, no differences between HIMA and PIMA were observed [5]. No differences were found at any intensities having the forearm positioned vertically [4]. Additionally, the averaged EMG at exhaustion as well as the power in the frequency range of 10 to 29 Hz were significantly higher during the force task compared to the position task. The EMG mean amplitude and mean frequency showed no significant differences between both tasks. [5] The recently published study [1] of our laboratory partly supported those results, partly disagreed with them. The setting included the recordings of mechanomyography (MMG) of the triceps brachii and the abdominal external oblique muscles as well as of mechanotendography (MTG) of the triceps brachii tendon, whereas no EMG was captured. The results revealed a significantly longer time to task failure (endurance time) at an intensity of 80% of the MVIC with vertically positioned forearm during PIMA compared to HIMA. The MMG amplitude of the triceps brachii in the last 10% of measurement time (exhaustion) was to be found significantly higher during PIMA vs. HIMA. The mean frequency did not differ significantly between both tasks. However, the MTG of triceps tendon showed a significantly higher power in the frequency ranges of 10 to 29 Hz and 8 to 15 Hz, respectively, during HIMA compared to PIMA. [1] This is opposite to the findings of Hunter et al. [7], who found a higher power during PIMA. The differences might be attributed to the different settings and, of course, different methods (EMG vs. MMG). Based on this, however, it is assumed that two different forms of isometric muscle action can be distinguished.

It is not this easy to enable the both forms of isometric muscle during measurements. In the former performed investigation, a pneumatic device was used so that the participant could realize the holding function [1]. Thereby, the push rod of the pneumatic system pushed against the forearm of the subject. Another possible approach to investigate the suspected two forms is that two persons interact mutually. Thereby, one partner pushes against the other one, while the second partner reacts to the force and resists by holding isometrically. This setting is assumed to be even more challenging, since two neuromuscular system interact. It could already be shown that in this kind of setting the MMG and MTG signals are able to develop coherent behavior in the sense of synchrony not only within one subject, but also between the muscles and tendons of both partners [8,9].

Using MMG, the mechanical micro-oscillations of muscles, which appear in frequency ranges around 10 Hz, are captured as one form of motor output [9,10]. This enables also to draw conclusions about central processes [11]. By maintaining a given force level, central structures must be activated to control the muscle action. Those motor control and regulating processes are assumed to get even more challenging, if the neuromuscular system has to react to exteroceptions. This is rather the case for the HIMA than for the PIMA.

The aim of the present study was to investigate whether or not the two forms of isometric muscle action can be distinguished by quantifiable parameters also during an interpersonal motor task.

## Methods

### Participants

In total, 20 healthy athletic subjects (*n* = 10 male, *n* = 10 female) volunteered to participate in the study, which were assigned to ten same sex couples for the measurements. The average age, weight and height of the male participants were 22.1 ± 2.4 years, 75.2 ± 6.9 kg and 181.5 ± 5.1 cm and 21.6 ± 2.1 years, 60.4 ± 3.5 kg and 168.3 ± 4.4 cm for the female participants. The averaged MVIC of the elbow extensors of the male participants amounted to 51.19 ± 22.45 Nm and the female participants reached a MVIC of averagely 25.85 ± 7.5 Nm (measurement position see below). Exclusion criteria were complaints of the upper extremities, shoulder girdle and spine within the las six month prior to measurements or any other health issue.

The study was approved by the local ethics committee at the University of Potsdam (Germany), approval no. 64/2016, and was conducted according to the Declaration of Helsinki. All participants were informed in detail and gave written informed consent to participate.

### Setting

The setting was identical to the investigations reported in Schaefer et al. [8] and Schaefer & Bittmann [9] (Fig. 1): The participants were sitting opposite. They were shifted slightly sidewise in a way, so that the measured dominant vertically positioned forearms were located in one plane. The angles between leg and trunk, arm and trunk as well as the elbow angle measured 90°. An interface with two shells of a thermic deformable polymer material (orthopedic material) was positioned proximal of the ulnar styloid processes of both participants and, thereby, connected the subjects. To record the reaction force between the participants, a strain gauge was fixed between the shells (model: ML MZ 2000 N 36, modified by biovision). Furthermore, a one-axial acceleration sensor (sensitivity: 312 mV/g, range ± 2 g, linearity: ± 0.2%; comp. biovision) was positioned on the strain gauge between the shells to capture the accelerations along the longitudinal acting force vector during the measurements. To record the mechanical musculo-tendinous oscillations of both participants (mechanomyography (MMG); mechanotendography (MTG)) piezoelectric sensors (Shadow SH 4001) were fixed using ECG-tape on the skin above the bellies of the lateral head of the triceps brachii muscle (MMGtri) and its tendon (MTGtri) as well as on the ipsilateral abdominal external oblique muscle (MMGobl; 2cm medial besides the free end of the 11^th^ rib). The triceps brachii muscle and its tendon were chosen since the main isometric muscle action in this position was performed by the elbow extensors. Thereby, the ipsilateral abdominal external oblique muscle served as an important stabilizer within the kinematic chain. The MMG-/MTG-signals were amplified by Nobels preamp booster pre-1, all signals were conducted to an A/D-converter (14-bit, National Instruments) and were recorded by the software NI DIAdem 10.2 (National Instruments) on a measurement notebook (Sony Vaio: PCG-61111M, Windows 7). The sampling rate was set at 1000 Hz. Between both subjects a “border” was constructed using a Thera band^®^. This was positioned in a way that, if the forearm of one participant touched the band, its elbow angle was extended by 10 degrees, thus, the partner had yielded by 10°. This was considered as leaving the isometric position and the measurement was stopped.

**Fig 1.**
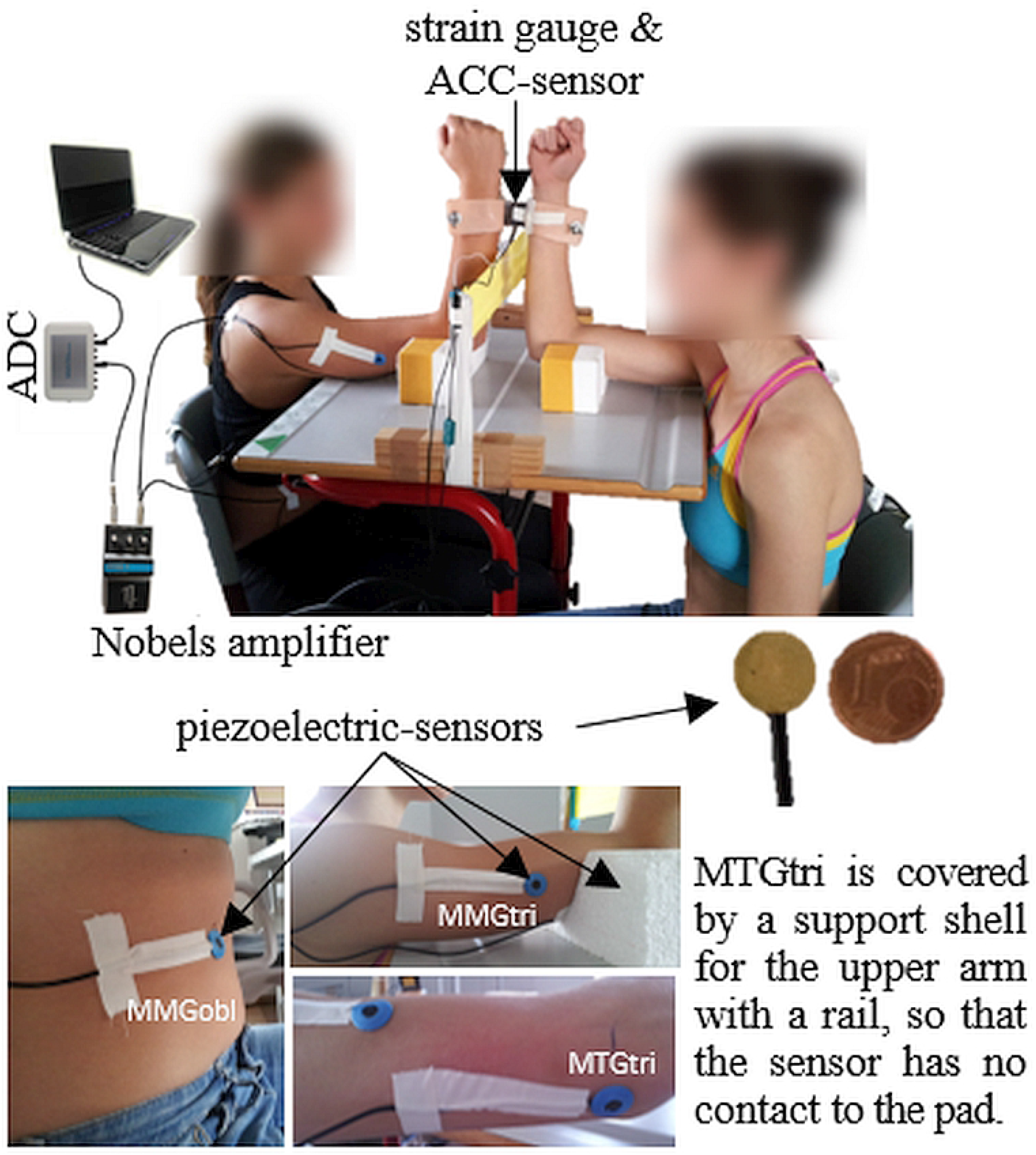
Setting. The partners are interacting with their distal forearms, which are coupled by two shells incl. the strain gauge and accelerometer. The piezoelectric sensor for MMG are fixed on the abdominal external oblique muscle (MMGobl), on the lateral head of the triceps brachii muscle (MMGtri) and – to capture the MTG of the triceps tendon (MTGtri) – on the olecranon fossa. The MMG/MTG-signals are amplified with the Nobels pre-amp booster pre-1. (Fig. 1 of Schaefer & Bittmann, 2018 [9]; modified with permission).

### Measuring tasks and procedure

The participants of each same sex pair should maintain an isometric muscle action using the elbow extensors at a reaction force of 80% of the MVIC (15s-trials) or 90% of the MVIC (fatiguing trials), respectively, of the weaker subject. The MVIC was recorded prior to the pair trials. In order to do this, each participant pushed singly in the measurement position against a shell of a fixed strain gauge, should reach its MVIC within 3s and should maintain it for 1s (2 trials). The position of the strain gauge was marked and the arm length was measured. The higher of both MVIC values of the weaker subject was used to calculate the force level of the pair trials. During the pair trials, each partner had different tasks, which changed within the measurement series: (1.) participant A pushed isometrically against participant B (PIMA), while (2.) participant B resisted the force applied by participant A isometrically in a holding manner (HIMA). Basically, PIMA is understood as an isometric muscle action in direction of elbow extension, like a kind of “stopped concentric action” without a change of joint angle. HIMA is seen as isometric reaction to an external force (here the force of the partner), which works in direction of elbow flexion. Thus, HIMA can be understood as a kind of “stopped eccentric action”. A more detailed description of HIMA and PIMA can be found in Schaefer & Bittmann.^1^ In the present setting, the holding participant should behave like a “wall” to hinder the partner to extend the arm, thus, maintaining the position ideally completely stable. In order to complete this task, the holding subject had to adapt to the applied force of the pushing partner. Therefore, the HIMA task required a high level of kinesthetic adaptation. If the holding partner yielded in the course of exhaustion, an elbow flexion was caused, including an eccentric muscle action of the triceps brachii muscle. The pushing partner controlled the force intensity via a dial instrument showed on a monitor in real time (biofeedback). Thus, the pushing partner had a visual feedback and should initiate the force. The isometric position can only be sustained if both partners maintain the given force level. As mentioned above, a deviation of 10° was interpreted as leaving the isometric position. This was the case, if one partner contacted the border (Thera band^®^) between the pair. Thereby, one partner flexed and the other partner extended the elbow joint with an amount of 10°. The pushing partner would contact the border, in case the holding counterpart would get tired first. Due to the task of the holding subject (just reacting to the force of the pushing partner), normally this subject should not be able to contact the border during the interaction. In case the pushing subject would get tired first, normally the holding partner should not extend his/her elbow angle, because he/she was instructed to stay in a stable position. In a case like this, the force level would decrease. Since this interaction is not trivial, the holding subject sometimes followed the yielding of the pushing partner in order to maintain the force level. Thereby, the initially pushing partner gave in and the formerly holding partner extended the elbow angle. This was the case in two trials of one single subject (see results and a critical discussion in the limitation section).

If the PIMA partner contacted the border or the HIMA partner aborted the interaction, it was classified as “HIMA failure”; if the PIMA partner get tired first and stopped the interaction or the HIMA partner contacted the border, respectively, the interaction was designated as “PIMA failure”. If both partners stopped the interaction simultaneously, the interaction was classified as even.

Two different measurement series were performed in the HIMA-PIMA-setting: (1) 15s-trials, whereby the isometric position should be sustained for 15s and (2) fatiguing trials, whereby the isometric position should be maintained as long as possible. The abortion criteria were either the contact to the border of one subject or a force decrease, whereby one or both partner stopped the interaction due to exhaustion.

In total, six 15s-trials were performed at 80% of the MVIC of the weaker person. In the first three of those six trials participant A performed PIMA and B performed HIMA, for the other three trials, the tasks changed (A: HIMA; B: PIMA). Afterwards, four fatiguing trials were performed at 90% of the MVIC. The higher intensity of 90% of the MVIC was chosen to limit the measurement duration. Again, the setting was complementary (PIMA vs. HIMA) with a change of the allocated task after the first two trials. The tasks for the participants to start with (HIMA or PIMA) were randomized for both measuring series. The resting period between the trials was 60s.

### Data processing

The main objective concerning the two forms of isometric muscle action is the endurance capability, thus, in the present setting who failed to maintain the task first compared between PIMA and HIMA. This was assessed qualitatively. If the PIMA partner contacted the border or if the HIMA partner quit the task, respectively, it was classified as “HIMA failure”. If the PIMA partner quit the task accompanied by a force break down, it was designated as “PIMA failure”. As mentioned above, one subject contacted the border during HIMA. This was categorized as “PIMA failure”, too. Nominal data were obtained (0 = “PIMA failure”, 1 = “HIMA failure”, 2 = “even”).

Considering the oscillating signals, the isometric plateau was cut from all signals (MMG, MTG) using the force signal. For that, the isometric plateau was defined as long as both partners were in contact. Slipping parts were excluded and in that case, the longer isometric plateau was used for further consideration. The signals of the isometric plateau were then filtered using a low pass Butterworth filter, (filter degree 5, cut-off frequency 20 Hz). Afterwards, the following parameters were considered: (1) the mean frequency, (2) the normalized mean amplitude, (3) the normalized amplitude variation, (4) the power in the frequency range of 8 to 15 Hz and (5) a special power-frequency ratio, which was recently used in another investigation concerning patients with Parkinson’s syndrome [12].

1. The mean frequency of the MMG-/MTG-signals was calculated using a Python script, which computed the average of the time intervals between two consecutive amplitude maxima and calculated the frequency thereof. According to Pikovsky et al. [13], this method is applicable for chaotic signals as neuromuscular ones.
2. The normalized mean amplitude of the MMG-/MTG-signals was evaluated in Excel (IBM Microsoft Office) by calculating the arithmetic mean of the amplitude maxima normalized to the maximal amplitude of the trial.
3. The normalized amplitude variation of the MMG-/MTG-signals was evaluated in Excel (IBM Microsoft Office). The differences between two consecutive maxima each were calculated and averaged over all time points. This mean variation value was normalized to the arithmetic mean of amplitude maxima of the trial.
4. The power spectral density (PSD) of the MMG-/MTG-signal was calculated in NI DIAdem 10.2. The values of power in the frequency range of 8 to 15 Hz was transferred to Excel, where the arithmetic mean of the power was calculated.
5. For the specific power frequency ratio MQ_REL_ the power spectral densities (PSD) of the MMG-/MTG-signals were calculated in NI DIAdem. Afterwards, the ratio of the power in the frequency range of 3 to 7 Hz to the sum of power in the frequency range of 3 to 7 and 7 to 12 Hz was calculated in Excel (IBM Microsoft Office) using the equation:

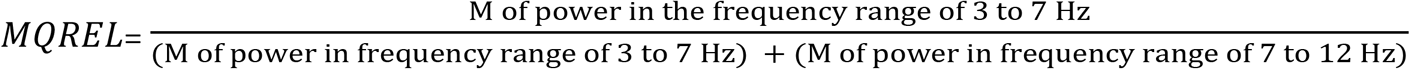

The arithmetic mean and coefficient of variation (CV) of all parameters (1) to (5) were calculated per participant, measurement series (15s-/fatiguing-trials) and tasks (HIMA vs. PIMA). The signals of one MMGtri, the MTGtri of one pair and one MMGobl signal were defective and therefore eliminated from the statistics.

#### Group statistics

IBM SPSS Statistics 26 was used for executing the group comparisons concerning the parameters (1) to (5) to investigate the differences between the motor tasks HIMA and PIMA. The significance level was set at *α* = 0.05. Firstly, the data were tested regarding their normal distribution by the Shapiro-Wilk-test. If the normal distribution was fulfilled, the paired t-test was utilized. The effect size was calculated by Pearson’s 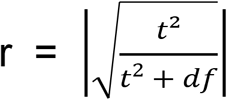. In case of non-parametric data, the comparison between HIMA and PIMA was done by means of the Wilcoxon-test for paired samples. The effect size was calculated by Pearson’s 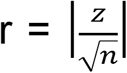. For comparing the nominal data (“HIMA failure” vs. “PIMA failure” vs. “even”) between HIMA and PIMA a χ^2^-test was performed in Excel (IBM Microsoft Office), the effect size was calculated by phi 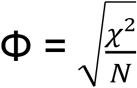.

## Results

Figure 2 displays exemplary raw signals of two fatiguing measurements of one pair at 90% of the MVIC during HIMA vs. PIMA.

**Fig 2.**
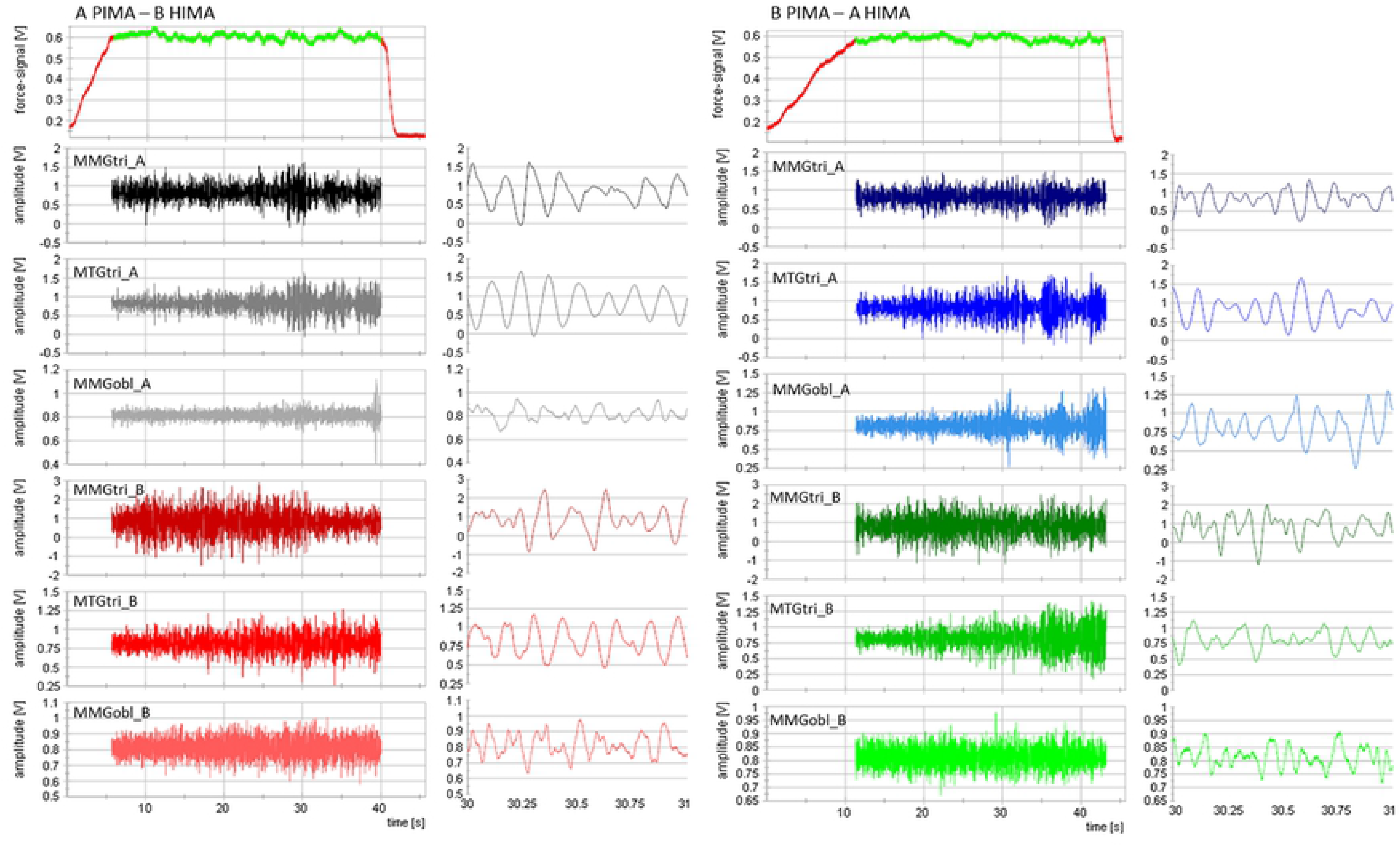
Exemplary raw signals. Displayed are the raw signals of two fatiguing measurements at 90% of the MVIC of one pair (A and B) of the strain gauge [V] and the mechanomyo-and -tendographic signals [V] of the triceps brachii and the abdominal external oblique muscles and of the triceps tendon, respectively. Left diagrams: A PIMA, B HIMA; right diagrams: A HIMA, B PIMA. The 1s-intervals to the right of each diagrams of the whole duration illustrate the good signal quality of the oscillatory raw signals.

### MVIC and force levels

The MVIC amounted averagely 51.19 ± 22.45 Nm for the male participants and 25.83 ± 0.07 Nm for the females. The averaged difference of MVIC values between the partners of one couple was 15.08 ± 9.40 Nm (24.56 ± 14.49%, range: 0.53 to 40.58%) for males and 3.53 ± 3.19 Nm (13.03 ± 11.09%, range: 1.2 to 27.33%) for females. Therefore, the adjusted force intensity of 80% of the MVIC of the weaker amounted for males averagely 62 ± 10% of the MVIC of the stronger partner; for the females it amounted 70 ± 9% for the stronger partner.

During the 15s-trials at 80% of the MVIC, the force level amounted averagely 23.08 ± 7.07 Nm during PIMA of subjects A (= HIMA of subjects B) and 23.12 ± 7.00 Nm during PIMA of subject B (= HIMA of subject A) (*W* = −0.563, *p* = 0.574). During the fatiguing trials at 90% of the MVIC, the force level amounted averagely 26.17 ± 8.08 Nm during PIMA of subjects A (= HIMA of subjects B) and 26.28 ± 7.90 Nm during PIMA of subject B (= HIMA of subject A) (*W* = 0.327, *p* = 0.744). Due to the non-significant differences between the tasks HIMA und PIMA, the consideration of the following other parameters are based on a similar force level.

### Capability of maintaining the tasks

The mean duration time of the fatiguing interactions over all trials was 24.58 ± 11.44s. Since in all interactions one partner performed PIMA and the other one performed HIMA, the duration time is not suitable to evaluate possible differences between HIMA and PIMA. That is why the endurance capability is captured qualitatively by counting the number of failures of each subject during the fatiguing trials. A “HIMA failure” occurred in 34 out of 40 trials (85%), whereas five PIMA failure arose (5 of 40; 12,5%). During one trial, the interaction was assessed as even, since both partners stopped the interaction simultaneously because of exhaustion (2.5%) (Fig. 3). The Chi-square test revealed a significant difference between PIMA and HIMA (*χ*^2^(2) = 25.87, *p* < 0.001) with a strong effect of Φ = 0.75.

**Fig 3.**
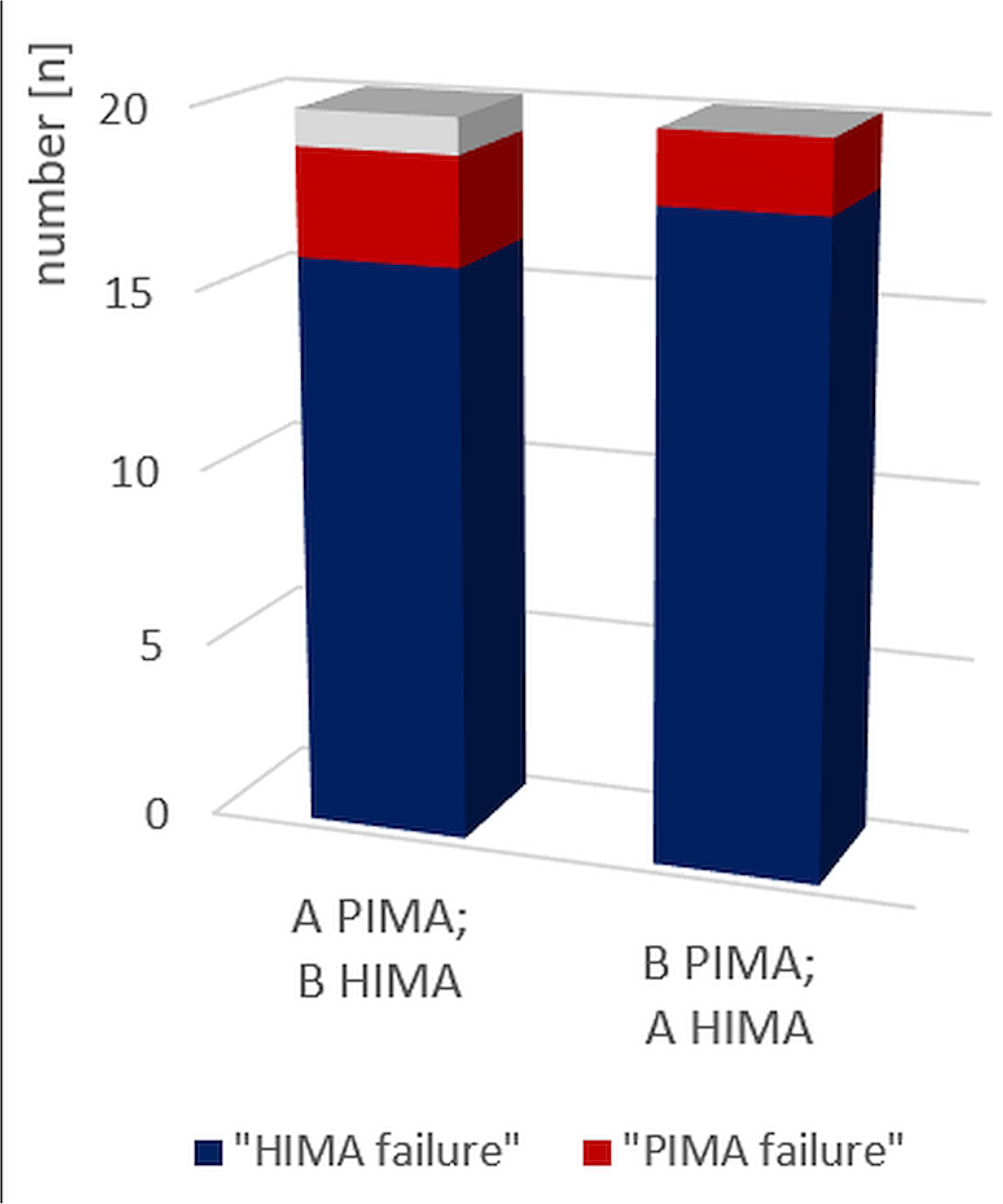
HIMA vs. PIMA failures. Displayed are the number of HIMA (n = 34, blue) and PIMA (n = 5, red) failures as well as the even (n = 1, grey) voted interaction during the 40 fatiguing trials (p < 0.001) grouped into subject A performs PIMA (B HIMA) vs. subject B performs PIMA (A HIMA).

### Amplitude and amplitude variation

The normalized amplitude showed no significant difference between HIMA and PIMA for MMGtri, MTGtri and MMGobl during the 15s and fatiguing trials. However, for MMGobl the comparison is just not significant, whereby during PIMA the normalized amplitude tends to have higher mean values compared to HIMA (*t*(19) = 2.048, *p* = 0.055, *r* = 0.42) (M and SD can be found in supplementary material).

The arithmetic means of variation of amplitude maxima revealed no significant difference between PIMA and HIMA for MMGtri and MTGtri, but they were significant for the MMGobl in the 15s-trials (t(19) = −2.759, *p* = 0.013, *r* = 0.55) and in the fatiguing trials (t(19) = −3.038, *p* = 0.007, *r* = 0.57) (Table 1, Fig. 4). Thereby, the amplitude variation was significantly higher during HIMA compared to PIMA. Looking at the single cases, 15 of 20 participants showed a higher amplitude variation during HIMA, although the couples did not change, only the tasks HIMA and PIMA.

**Table 1.**
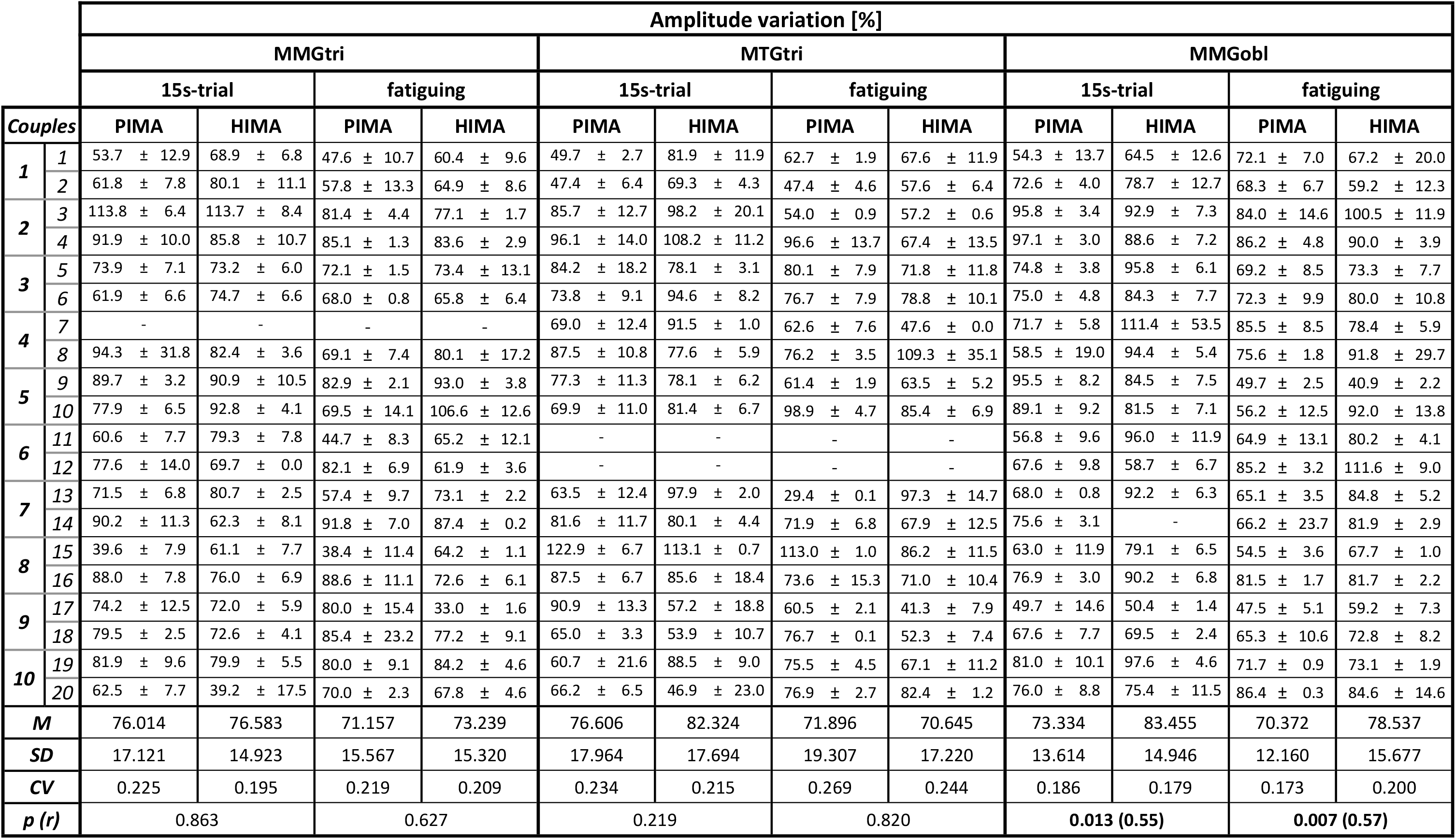
Amplitude variation. Arithmetic means (M) (± standard deviation (SD)) of the amplitude variation [%] of the mechanomyographic and mechanotendographic signals of the triceps brachii muscle (MMGtri) and its tendon (MTGtri) as well as of the abdominal external oblique muscle (MMGobl) during the 15s and fatiguing trials comparing PIMA vs. HIMA. The group M, SD, coefficient of variation (CV) and p-values of statistical comparisons between HIMA and PIMA are displayed. In case of significance, the effect size r is given.

**Fig 4.**
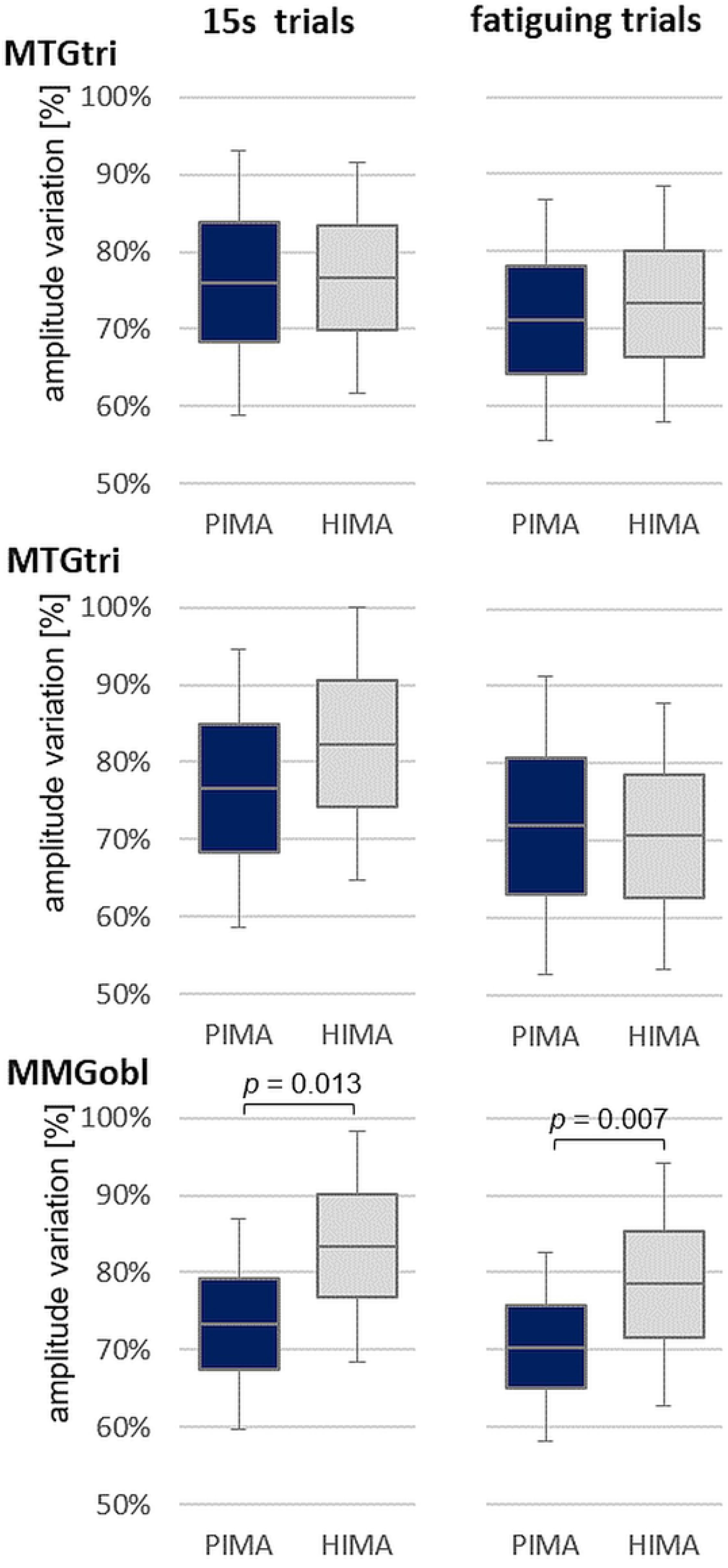
Amplitude variation. Displayed are the arithmetic means, standard deviations and the 95%-confidence intervals of the amplitude variation [%] of the signals MMGtri, MTGtri and MMGobl during the 15s trials (left) and fatiguing trials (right) comparing PIMA (blue) vs. HIMA (grey). T-test was significant for MMGobl.

### Frequency and power

During the 15s and fatiguing trials, the arithmetic mean of mean frequency showed no significant differences in the paired t-test between PIMA and HIMA (*p* > 0.05). The CV of frequency between the fatiguing trials was significantly higher during HIMA compared to PIMA, however, this only occurred for the MMGtri (*W* = 2.575, *p* = 0.010, *r* = 0.59). (M and SD can be found in the supplementary material)

Considering the frequency range of 8 to 15 Hz the power was significantly different for MMGobl during the 15s-trials (*W* = −3.300, *p* = 0.001, *r* = 0.76) and during the fatiguing trials (*W* = −2.539, *p* = 0.011, *r* = 0.57), whereby PIMA showed higher values than HIMA (see Table 2, Fig. 5). MMGtri and MTGtri showed no significances thereby (*p* > 0.05).

**Table 2.** Power in 8 to 15 Hz. Arithmetic means (M) (± standard deviation (SD)) of the power in the frequency range of 8 to 15 Hz [V²/Hz] of the mechanomyographic and mechanotendographic signals of the triceps brachii muscle (MMGtri) and its tendon (MTGtri) as well as of the abdominal external oblique muscle (MMGobl) during the 15s and fatiguing trials comparing PIMA vs. HIMA. The group M, SD, coefficient of variation (CV) and p-values of statistical comparisons between HIMA and PIMA are displayed. In case of significance, the effect size r is given.

| Power in frequency range of 8 to 15 Hz [$V^2/Hz$] | | | | | | | | | | | | | |
| --- | --- | --- | --- | --- | --- | --- | --- | --- | --- | --- | --- | --- | --- |
|  |  | MMGtri |  |  |  | MTGtri |  |  |  | MMGobl |  |  |  |
|  |  | 15s-trial |  | fatiguing |  | 15s-trial |  | fatiguing |  | 15s-trial |  | fatiguing |  |
| Couples |  | PIMA | HIMA | PIMA | HIMA | PIMA | HIMA | PIMA | HIMA | PIMA | HIMA | PIMA | HIMA |
| 1 | 1 | 0.026 ± 0.014 | 0.032 ± 0.016 | 0.150 ± 0.029 | 0.127 ± 0.046 | 0.051 ± 0.018 | 0.018 ± 0.003 | 0.014 ± 0.012 | 0.022 ± 0.003 | 0.008 ± 0.003 | 0.004 ± 0.001 | 0.010 ± 0.001 | 0.013 ± 0.012 |
| | 2 | 0.035 ± 0.008 | 0.016 ± 0.006 | 0.042 ± 0.006 | 0.059 ± 0.032 | 0.022 ± 0.008 | 0.005 ± 0.002 | 0.074 ± 0.006 | 0.028 ± 0.006 | 0.000 ± 0.000 | 0.000 ± 0.000 | 0.003 ± 0.001 | 0.003 ± 0.000 |
| 2 | 3 | 0.001 ± 0.001 | 0.000 ± 0.000 | 0.004 ± 0.000 | 0.006 ± 0.006 | 0.000 ± 0.000 | 0.000 ± 0.000 | 0.004 ± 0.004 | 0.002 ± 0.002 | 0.000 ± 0.000 | 0.000 ± 0.000 | 0.001 ± 0.001 | 0.000 ± 0.000 |
| | 4 | 0.003 ± 0.001 | 0.009 ± 0.004 | 0.028 ± 0.002 | 0.023 ± 0.002 | 0.001 ± 0.000 | 0.000 ± 0.000 | 0.001 ± 0.000 | 0.002 ± 0.002 | 0.000 ± 0.000 | 0.000 ± 0.000 | 0.000 ± 0.000 | 0.000 ± 0.000 |
| 3 | 5 | 0.001 ± 0.000 | 0.001 ± 0.000 | 0.004 ± 0.002 | 0.007 ± 0.000 | 0.000 ± 0.000 | 0.000 ± 0.000 | 0.000 ± 0.000 | 0.002 ± 0.002 | 0.000 ± 0.000 | 0.000 ± 0.000 | 0.000 ± 0.000 | 0.000 ± 0.000 |
| | 6 | 0.003 ± 0.002 | 0.000 ± 0.000 | 0.015 ± 0.011 | 0.006 ± 0.002 | 0.000 ± 0.000 | 0.000 ± 0.000 | 0.002 ± 0.000 | 0.000 ± 0.000 | 0.000 ± 0.000 | 0.000 ± 0.000 | 0.001 ± 0.000 | 0.000 ± 0.000 |
| 4 | 7 | - | - | - | - | 0.029 ± 0.012 | 0.010 ± 0.005 | 0.012 ± 0.008 | 0.024 ± | 0.000 ± 0.000 | 0.001 ± 0.001 | 0.000 ± 0.000 | 0.000 ± 0.000 |
| | 8 | 0.007 ± 0.006 | 0.006 ± 0.002 | 0.011 ± 0.003 | 0.002 ± 0.003 | 0.074 ± 0.082 | 0.059 ± 0.041 | 0.052 ± 0.071 | 0.014 ± 0.007 | 0.005 ± 0.005 | 0.002 ± 0.001 | 0.002 ± 0.002 | 0.001 ± 0.001 |
| 5 | 9 | 0.001 ± 0.000 | 0.002 ± 0.000 | 0.003 ± 0.001 | 0.004 ± 0.001 | 0.003 ± 0.001 | 0.002 ± 0.001 | 0.005 ± 0.001 | 0.011 ± 0.000 | 0.000 ± 0.000 | 0.000 ± 0.000 | 0.000 ± 0.000 | 0.001 ± 0.000 |
| | 10 | 0.001 ± 0.001 | 0.001 ± 0.000 | 0.003 ± 0.002 | 0.001 ± 0.000 | 0.003 ± 0.001 | 0.002 ± 0.000 | 0.001 ± 0.000 | 0.001 ± 0.000 | 0.000 ± 0.000 | 0.000 ± 0.000 | 0.004 ± 0.003 | 0.001 ± 0.000 |
| 6 | 11 | 0.129 ± 0.053 | 0.047 ± 0.005 | 0.170 ± 0.087 | 0.075 ± 0.044 | - | - | - | - | 0.005 ± 0.001 | 0.000 ± 0.000 | 0.005 ± 0.002 | 0.001 ± 0.001 |
| | 12 | 0.060 ± 0.048 | 0.089 ± 0.010 | 0.084 ± 0.035 | 0.098 ± 0.048 | - | - | - | - | 0.001 ± 0.000 | 0.000 ± 0.000 | 0.000 ± 0.000 | 0.000 ± 0.000 |
| 7 | 13 | 0.005 ± 0.001 | 0.003 ± 0.001 | 0.021 ± 0.003 | 0.002 ± 0.000 | 0.008 ± 0.003 | 0.001 ± 0.000 | 0.038 ± 0.004 | 0.001 ± 0.000 | 0.007 ± 0.001 | 0.002 ± 0.001 | 0.013 ± 0.002 | 0.007 ± 0.002 |
| | 14 | 0.005 ± 0.002 | 0.003 ± 0.001 | 0.009 ± 0.002 | 0.012 ± 0.002 | 0.007 ± 0.001 | 0.008 ± 0.001 | 0.017 ± 0.014 | 0.023 ± 0.011 | 0.001 ± 0.000 | - | 0.004 ± 0.002 | 0.003 ± 0.000 |
| 8 | 15 | 0.010 ± 0.005 | 0.003 ± 0.001 | 0.016 ± 0.008 | 0.006 ± 0.001 | 0.000 ± 0.000 | 0.000 ± 0.000 | 0.001 ± 0.000 | 0.002 ± 0.001 | 0.001 ± 0.000 | 0.000 ± 0.000 | 0.002 ± 0.000 | 0.001 ± 0.000 |
| | 16 | 0.004 ± 0.001 | 0.003 ± 0.001 | 0.006 ± 0.000 | 0.008 ± 0.006 | 0.004 ± 0.001 | 0.002 ± 0.001 | 0.011 ± 0.008 | 0.018 ± 0.018 | 0.001 ± 0.000 | 0.000 ± 0.000 | 0.001 ± 0.000 | 0.001 ± 0.001 |
| 9 | 17 | 0.005 ± 0.002 | 0.007 ± 0.004 | 0.013 ± 0.008 | 0.038 ± 0.019 | 0.005 ± 0.001 | 0.019 ± 0.008 | 0.029 ± 0.004 | 0.031 ± 0.022 | 0.000 ± 0.000 | 0.000 ± 0.000 | 0.000 ± 0.000 | 0.003 ± 0.001 |
| | 18 | 0.031 ± 0.008 | 0.016 ± 0.004 | 0.036 ± 0.045 | 0.016 ± 0.000 | 0.052 ± 0.050 | 0.023 ± 0.007 | 0.008 ± 0.002 | 0.008 ± 0.004 | 0.003 ± 0.001 | 0.001 ± 0.000 | 0.005 ± 0.002 | 0.002 ± 0.001 |
| 10 | 19 | 0.038 ± 0.003 | 0.052 ± 0.006 | 0.051 ± 0.006 | 0.051 ± 0.005 | 0.011 ± 0.003 | 0.004 ± 0.001 | 0.006 ± 0.000 | 0.003 ± 0.001 | 0.000 ± 0.000 | 0.000 ± 0.000 | 0.001 ± 0.000 | 0.001 ± 0.000 |
| | 20 | 0.127 ± 0.033 | 0.252 ± 0.049 | 0.066 ± 0.023 | 0.116 ± 0.008 | 0.007 ± 0.002 | 0.023 ± 0.016 | 0.005 ± 0.000 | 0.011 ± 0.003 | 0.007 ± 0.002 | 0.006 ± 0.000 | 0.012 ± 0.001 | 0.008 ± 0.004 |
| **M** |  | 0.026 | 0.029 | 0.039 | 0.035 | 0.015 | 0.010 | 0.016 | 0.011 | 0.002 | 0.001 | 0.003 | 0.002 |
| **SD** |  | 0.040 | 0.059 | 0.049 | 0.041 | 0.022 | 0.015 | 0.020 | 0.011 | 0.003 | 0.002 | 0.004 | 0.003 |
| **CV** |  | 1.533 | 2.071 | 1.262 | 1.198 | 1.430 | 1.508 | 1.304 | 0.932 | 1.373 | 1.744 | 1.228 | 1.422 |
| **p (r)** |  | 0.717 |  | 0.601 |  | 0.053 |  | 0.679 |  | 0.001 (0.76) |  | 0.011 (0.57) |  |

**Fig 5.**
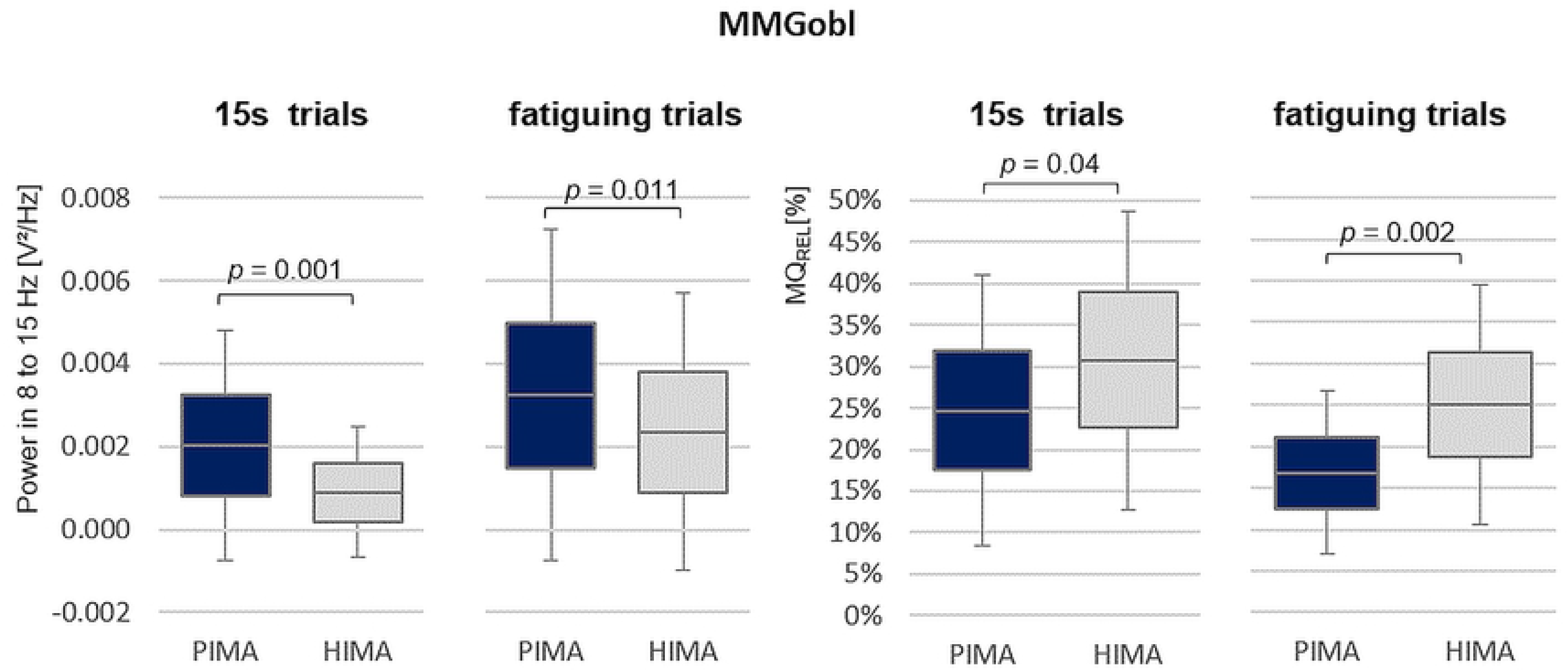
Power in 8 to 15 Hz and power frequency ratio MQ_REL_. Displayed are the arithmetic means, standard deviations and the 95%-confidence intervals of the power in the frequency range of 8 to 15 Hz (left) and of the power frequency ratio MQ_REL_ of the MMGobl signal during the 15s and fatiguing trials for the tasks PIMA (blue) and HIMA (grey). Statistical comparisons between PIMA and HIMA revealed significant results for all comparisons with p = 0.001 to 0.04 and effect sizes of r = 0.47 to 0.76.

Furthermore, the power-frequency-ratio MQ_REL_ showed significantly higher values in the MMGobl-signal during HIMA compared to PIMA during the fatiguing trials (t(19) = −3.673, *p* = 0.002, *r* = 0.64) and during the 15s-trials (*W* = 2.052, *p* = 0.040, *r* = 0.47) (Table 3, Fig. 5). Thereby, during HIMA, the lower frequency range of 3 to 7 Hz amounted averagely 24.9 ± 14.6% of the whole range of 3 to 12 Hz during the fatiguing trials and averagely 30.8 ± 18.0% during 15s-trials. Whereas during PIMA, the lower frequency range amounted 16.7 ± 9.9% of the whole frequency range of 3 to 12 Hz during the fatiguing trials and 24.7 ± 16.3 % during the 15s-trials. The signals of MMGtri and MTGtri did not differ significantly concerning those parameters (*p* > 0.05) (see Table 3).

**Table 3.**
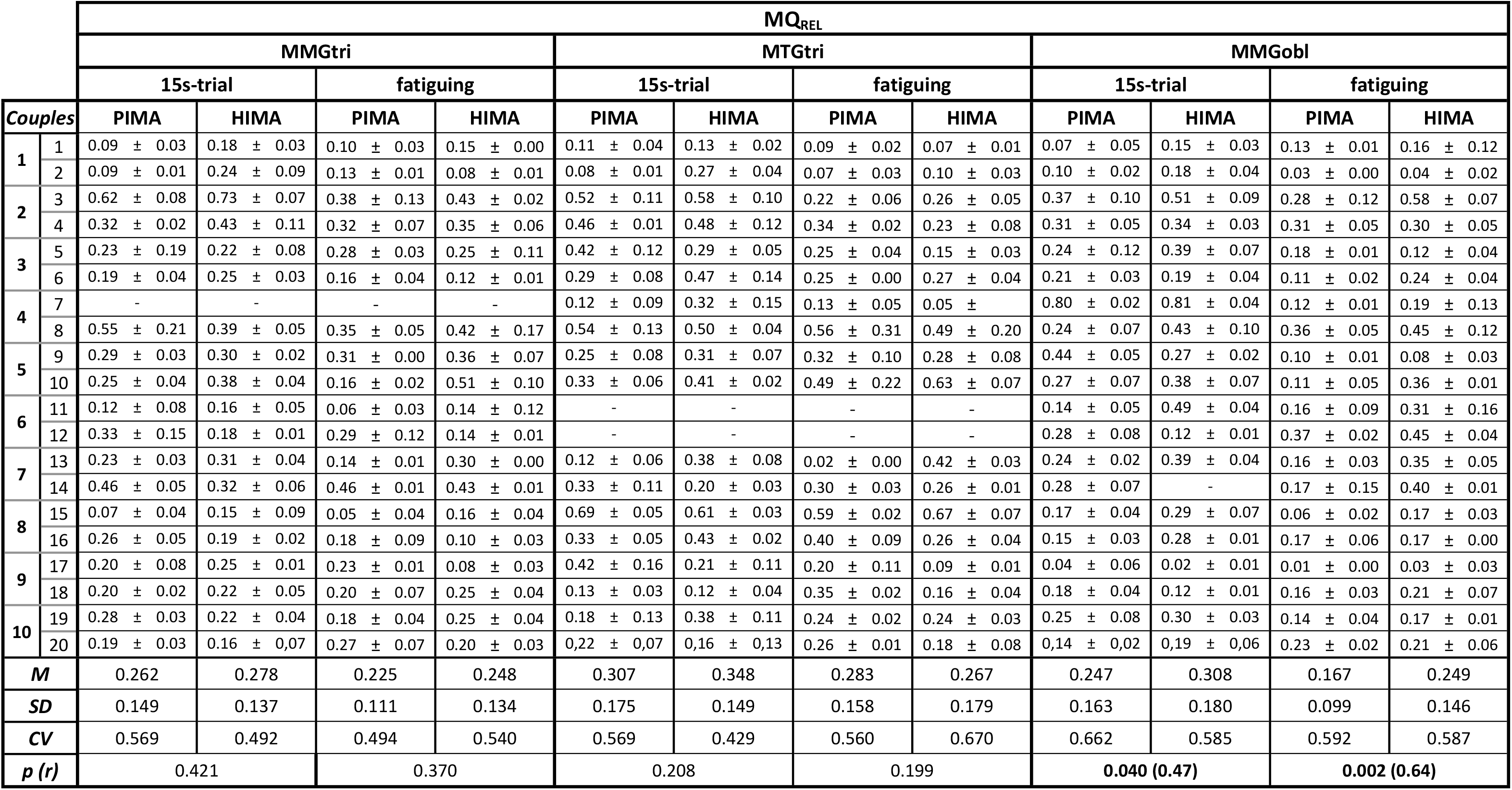
Power-frequency ratio MQ_REL_. Arithmetic means (M) (± standard deviation (SD)) of the power-frequency ratio MQ_REL_ of the mechanomyographic and mechanotendographic signals of the triceps brachii muscle (MMGtri) and its tendon (MTGtri) as well as of the abdominal external oblique muscle (MMGobl) during the 15s and fatiguing trials comparing PIMA vs. HIMA. The group means, SD, coefficient of variation (CV) and p-values of statistical comparisons between HIMA and PIMA are displayed. In case of significance, the effect size r is given.

| $MQ_{REL}$ | | | | | | | | | | | | | |
| --- | --- | --- | --- | --- | --- | --- | --- | --- | --- | --- | --- | --- | --- |
|  |  | MMGtri |  |  |  | MTGtri |  |  |  | MMGobl |  |  |  |
|  |  | 15s-trial |  | fatiguing |  | 15s-trial |  | fatiguing |  | 15s-trial |  | fatiguing |  |
| Couples |  | PIMA | HIMA | PIMA | HIMA | PIMA | HIMA | PIMA | HIMA | PIMA | HIMA | PIMA | HIMA |
| 1 | 1 | 0.09 ± 0.03 | 0.18 ± 0.03 | 0.10 ± 0.03 | 0.15 ± 0.00 | 0.11 ± 0.04 | 0.13 ± 0.02 | 0.09 ± 0.02 | 0.07 ± 0.01 | 0.07 ± 0.05 | 0.15 ± 0.03 | 0.13 ± 0.01 | 0.16 ± 0.12 |
| | 2 | 0.09 ± 0.01 | 0.24 ± 0.09 | 0.13 ± 0.01 | 0.08 ± 0.01 | 0.08 ± 0.01 | 0.27 ± 0.04 | 0.07 ± 0.03 | 0.10 ± 0.03 | 0.10 ± 0.02 | 0.18 ± 0.04 | 0.03 ± 0.00 | 0.04 ± 0.02 |
| 2 | 3 | 0.62 ± 0.08 | 0.73 ± 0.07 | 0.38 ± 0.13 | 0.43 ± 0.02 | 0.52 ± 0.11 | 0.58 ± 0.10 | 0.22 ± 0.06 | 0.26 ± 0.05 | 0.37 ± 0.10 | 0.51 ± 0.09 | 0.28 ± 0.12 | 0.58 ± 0.07 |
| | 4 | 0.32 ± 0.02 | 0.43 ± 0.11 | 0.32 ± 0.07 | 0.35 ± 0.06 | 0.46 ± 0.01 | 0.48 ± 0.12 | 0.34 ± 0.02 | 0.23 ± 0.08 | 0.31 ± 0.05 | 0.34 ± 0.03 | 0.31 ± 0.05 | 0.30 ± 0.05 |
| 3 | 5 | 0.23 ± 0.19 | 0.22 ± 0.08 | 0.28 ± 0.03 | 0.25 ± 0.11 | 0.42 ± 0.12 | 0.29 ± 0.05 | 0.25 ± 0.04 | 0.15 ± 0.03 | 0.24 ± 0.12 | 0.39 ± 0.07 | 0.18 ± 0.01 | 0.12 ± 0.04 |
| | 6 | 0.19 ± 0.04 | 0.25 ± 0.03 | 0.16 ± 0.04 | 0.12 ± 0.01 | 0.29 ± 0.08 | 0.47 ± 0.14 | 0.25 ± 0.00 | 0.27 ± 0.04 | 0.21 ± 0.03 | 0.19 ± 0.04 | 0.11 ± 0.02 | 0.24 ± 0.04 |
| 4 | 7 | - | - | - | - | 0.12 ± 0.09 | 0.32 ± 0.15 | 0.13 ± 0.05 | 0.05 ± | 0.80 ± 0.02 | 0.81 ± 0.04 | 0.12 ± 0.01 | 0.19 ± 0.13 |
| | 8 | 0.55 ± 0.21 | 0.39 ± 0.05 | 0.35 ± 0.05 | 0.42 ± 0.17 | 0.54 ± 0.13 | 0.50 ± 0.04 | 0.56 ± 0.31 | 0.49 ± 0.20 | 0.24 ± 0.07 | 0.43 ± 0.10 | 0.36 ± 0.05 | 0.45 ± 0.12 |
| 5 | 9 | 0.29 ± 0.03 | 0.30 ± 0.02 | 0.31 ± 0.00 | 0.36 ± 0.07 | 0.25 ± 0.08 | 0.31 ± 0.07 | 0.32 ± 0.10 | 0.28 ± 0.08 | 0.44 ± 0.05 | 0.27 ± 0.02 | 0.10 ± 0.01 | 0.08 ± 0.03 |
| | 10 | 0.25 ± 0.04 | 0.38 ± 0.04 | 0.16 ± 0.02 | 0.51 ± 0.10 | 0.33 ± 0.06 | 0.41 ± 0.02 | 0.49 ± 0.22 | 0.63 ± 0.07 | 0.27 ± 0.07 | 0.38 ± 0.07 | 0.11 ± 0.05 | 0.36 ± 0.01 |
| 6 | 11 | 0.12 ± 0.08 | 0.16 ± 0.05 | 0.06 ± 0.03 | 0.14 ± 0.12 | - | - | - | - | 0.14 ± 0.05 | 0.49 ± 0.04 | 0.16 ± 0.09 | 0.31 ± 0.16 |
| | 12 | 0.33 ± 0.15 | 0.18 ± 0.01 | 0.29 ± 0.12 | 0.14 ± 0.01 | - | - | - | - | 0.28 ± 0.08 | 0.12 ± 0.01 | 0.37 ± 0.02 | 0.45 ± 0.04 |
| 7 | 13 | 0.23 ± 0.03 | 0.31 ± 0.04 | 0.14 ± 0.01 | 0.30 ± 0.00 | 0.12 ± 0.06 | 0.38 ± 0.08 | 0.02 ± 0.00 | 0.42 ± 0.03 | 0.24 ± 0.02 | 0.39 ± 0.04 | 0.16 ± 0.03 | 0.35 ± 0.05 |
| | 14 | 0.46 ± 0.05 | 0.32 ± 0.06 | 0.46 ± 0.01 | 0.43 ± 0.01 | 0.33 ± 0.11 | 0.20 ± 0.03 | 0.30 ± 0.03 | 0.26 ± 0.01 | 0.28 ± 0.07 | - | 0.17 ± 0.15 | 0.40 ± 0.01 |
| 8 | 15 | 0.07 ± 0.04 | 0.15 ± 0.09 | 0.05 ± 0.04 | 0.16 ± 0.04 | 0.69 ± 0.05 | 0.61 ± 0.03 | 0.59 ± 0.02 | 0.67 ± 0.07 | 0.17 ± 0.04 | 0.29 ± 0.07 | 0.06 ± 0.02 | 0.17 ± 0.03 |
| | 16 | 0.26 ± 0.05 | 0.19 ± 0.02 | 0.18 ± 0.09 | 0.10 ± 0.03 | 0.33 ± 0.05 | 0.43 ± 0.02 | 0.40 ± 0.09 | 0.26 ± 0.04 | 0.15 ± 0.03 | 0.28 ± 0.01 | 0.17 ± 0.06 | 0.17 ± 0.00 |
| 9 | 17 | 0.20 ± 0.08 | 0.25 ± 0.01 | 0.23 ± 0.01 | 0.08 ± 0.03 | 0.42 ± 0.16 | 0.21 ± 0.11 | 0.20 ± 0.11 | 0.09 ± 0.01 | 0.04 ± 0.06 | 0.02 ± 0.01 | 0.01 ± 0.00 | 0.03 ± 0.03 |
| | 18 | 0.20 ± 0.02 | 0.22 ± 0.05 | 0.20 ± 0.07 | 0.25 ± 0.04 | 0.13 ± 0.03 | 0.12 ± 0.04 | 0.35 ± 0.02 | 0.16 ± 0.04 | 0.18 ± 0.04 | 0.12 ± 0.01 | 0.16 ± 0.03 | 0.21 ± 0.07 |
| 10 | 19 | 0.28 ± 0.03 | 0.22 ± 0.04 | 0.18 ± 0.04 | 0.25 ± 0.04 | 0.18 ± 0.13 | 0.38 ± 0.11 | 0.24 ± 0.02 | 0.24 ± 0.03 | 0.25 ± 0.08 | 0.30 ± 0.03 | 0.14 ± 0.04 | 0.17 ± 0.01 |
| | 20 | 0.19 ± 0.03 | 0.16 ± 0.07 | 0.27 ± 0.07 | 0.20 ± 0.03 | 0.22 ± 0.07 | 0.16 ± 0.13 | 0.26 ± 0.01 | 0.18 ± 0.08 | 0.14 ± 0.02 | 0.19 ± 0.06 | 0.23 ± 0.02 | 0.21 ± 0.06 |
| **M** |  | 0.262 | 0.278 | 0.225 | 0.248 | 0.307 | 0.348 | 0.283 | 0.267 | 0.247 | 0.308 | 0.167 | 0.249 |
| **SD** |  | 0.149 | 0.137 | 0.111 | 0.134 | 0.175 | 0.149 | 0.158 | 0.179 | 0.163 | 0.180 | 0.099 | 0.146 |
| **CV** |  | 0.569 | 0.492 | 0.494 | 0.540 | 0.569 | 0.429 | 0.560 | 0.670 | 0.662 | 0.585 | 0.592 | 0.587 |
| **p (r)** |  | 0.421 |  | 0.370 |  | 0.208 |  | 0.199 |  | 0.040 (0.47) |  | 0.002 (0.64) |  |

## Discussion

In the present study, two tasks of isometric muscle action were performed in a paired-couple-setting, whereby each of the two partners performed either the pushing isometric motor task (PIMA) or the holding isometric motor task (HIMA). The tasks changed, so that each partner performed each task in the same couple. The main findings were: (1) The subject performing PIMA was able to maintain the force level for longer, the categorization “HIMA failure” arised in 85% of the 40 fatiguing trials (*p* < 0.000). (2) The mean frequency showed no significant differences between PIMA and HIMA for all signals in 15s and fatiguing trials (p > 0.05). (3) Concerning the normalized amplitude only the MMGobl tend to have higher values during PIMA than HIMA (*p* = 0.055, n.s.). (4) MMGtri and MTGtri showed no significant differences between PIMA and HIMA regarding the other oscillatory parameters (amplitude variation, power and MQ_REL_). The MMGobl showed significant differences between PIMA and HIMA during the 15s and fatiguing trials concerning (5) the amplitude variation (15s: *p* = 0.013; *r* = 0.55, fatigue: *p* = 0.007, *r* = 0.57), whereby the amplitude varies more during HIMA; (6) the power in the frequency range of 8 to 15 Hz (*p* = 0.001, *r* = 0.76; *p* = 0.011, *r* = 0.57), whereby the power is higher during PIMA and concerning (7) the power-frequency ratio MQ_REL_ *(p* = 0.040, *r* = 0.47; *p* = 0.002, *r* = 0.64), whereby during HIMA the power in the frequency range of 3 to 7 Hz related to the frequency range of 3 to 12 Hz is higher compared to PIMA.

### Methodical limitations

The present setting of muscular interaction, whereby one partner performed HIMA and the other one performed PIMA, is a novel approach. Thereby, the inspection of the performed motor tasks is difficult. This limitation was tried to be controlled by the design that only the pushing partner had the visual control of the force level and, therefore, should act and initiate the force isometrically in direction of elbow extension. The holding partner should only react isometrically and, thereby, just had a kinesthetic control. Therefore, the task instructions were given in a pronounced way to ensure the correct performances of both tasks. However, it could not be ruled out completely that a subject switched unconsciously between the tasks HIMA and PIMA in the course of measurements.

As mentioned above, one participant contacted the border during HIMA. This was not expected since, thereby, the subject extended its elbow joint although this is not intended during HIMA. That is why we assume that this participant switched into PIMA in the course of the interaction. The other partner yielded and, thus, flexed his elbow. Thus, an eccentric muscle action must have been performed. This should not occur during PIMA – per definitionem.

Since this was the only exception, we assume that the tasks were performed largely adequate due to the following reasons: the tasks instructions, the setting (control of force level only by the pushing subject, the holding one just reacted to the partners force) and the fact that the border was contacted in most cases by the pushing partner – irrelevant of the physical condition of the partners.

Another limit was that during the interaction, the partners sometimes slipped, so that the measurement had to be repeated. This was due to the connection with the strain gauge, the oscillating character of the interaction and the high force level. For further investigations, the force level might be reduced, so that the problem would be solved.

Furthermore, due to the different physical conditions, the force level was not always similar for both partners. This is a limitation, which is difficult to solve. However, since the tasks changed, this limit was controlled. Nevertheless, the results that the pushing partner mostly could maintain the task – irrespective of the physical condition – is even more impressive due to the different physical conditions.

For further investigations, the MVIC could be measured during interaction between the partners, whereby both partners perform PIMA, so that the maximal reaction force of the pair would be regarded as MVIC. There are pros and cons for both alternatives.

Four MMG/MTG-signals had to be excluded, since the sensors failed. This could have been controlled better in the course of the measurements. However, due to the high number of MMG- and MTG-sensors, there was no opportunity to exchange the piezoelectric sensors during the measuring.

A multiple test problem could arise for critical statisticians. Since the study has an explorative character, this limitation is discussible. Many researchers take the view that multiple testing is permitted in this case [14–17]. To gather an initial suspicion, there is the need of an extensive consideration to look into several possibilities. However, the results have to be interpreted with caution – also because the small sample size of 10 pairs.

### Capability of maintaining the tasks PIMA or HIMA

It is noteworthy that the results were obtained in a setting, in which the tasks changed in the same couple. Thus, depending on the tasks HIMA or PIMA, the capacity of maintaining the identical force level changed – irrelevant, which partner was physically stronger and showed a higher MVIC. This was also apparent for couples with very high differences in MVIC, e.g. in one couple the stronger partner had a MVIC of 159% of the MVIC of the weaker. That means he performed the fatiguing trials with 53% of his MVIC. Nevertheless, he failed during the HIMA trials. 34 “HIMA failures” out of 40 trials and, thus, a better capability of maintaining the task during PIMA suggest a better submaximal isometric endurance during a pushing isometric action compared to a holding one with identical force. Because of the randomized order of the allocation of the tasks, an effect of the physical conditions can be ruled out as reason for the results of longer maintaining time in PIMA. Only five “PIMA failures” occurred, whereby just one subject contacted the border in two trials. We assume that the holding partner switched into PIMA in the course of interaction. It was obvious that in those trials the pushing partner started to move back and the formerly holding partner followed him and extended his elbow angle in that process – so probably switched into PIMA. For what reason this outcome appeared can only be discussed hypothetically. The MVIC was nearly identical between both partners (partner A: 33.18 Nm vs. B: 32.90 Nm).

There are different possible explanations for the decreased capability to maintain the HIMA task, e.g. a metabolic fatigue, effects of ischemia or higher demands of neural control.

Basically, a metabolic fatigue seems to be less obvious, since the intensity and muscle length during the isometric muscle action were similar for both tasks. Nevertheless, a possible higher demand of energetic substances or a higher production of lactate during HIMA would be conceivable. In a study of Rudroff et al. [6] a higher glucose uptake (PET method) of the lower extremity muscles was found in 11 of 24 muscles after fatiguing trials at 25% of MVIC performing HIMA compared to PIMA in young male (*n* = 3), but not for older male participants (*n* = 3). This would speak for a higher consumption of glucose during HIMA in young men. Another explanation for the shorter endurance time during HIMA could be the capillary behavior. There seems to be different types of capillary behavior as shown in Dech et al. [18] during holding a weight. The different isometric tasks were not considered thereby. However, it cannot be ruled out that different behaviors during HIMA and PIMA might exist. Nevertheless, in a study of Booghs et al. [19] no difference in muscle oxygenation of the elbow flexors at 20 and 60% of the MVIC occurred comparing HIMA vs. PIMA. Since the muscle oxygenation was not considered in the presented study, no statement can be made concerning this parameter. Furthermore, a central fatigue due to higher neuronal demands could also be assumed for the briefer endurance time during HIMA.

### Possible higher demands of neuronal control during HIMA compared to PIMA

It was supposed that HIMA requires higher neural control strategies compared to PIMA [1], since during HIMA the endurance time was significantly shorter compared to PIMA in several investigations and in the present study, too [1,4–7]. In the present study, the myofascial oscillations of the investigated muscles showed no differences between HIMA and PIMA concerning the mean frequency and mean amplitude of all signals. This was hypothesized on the base of the former MMG/MTG study during single measurements [1] as well as on other EMG studies [2–5] investigating HIMA and PIMA. The commonly considered parameters of frequency and amplitude of EMG or MMG/MTG, respectively, seem not to provide a suitable approach to distinguish HIMA and PIMA. This underlines the necessity to consider other parameters, provided that there is a difference in performing those tasks, which is reflected by the maintaining time.

Assuming different neural control strategies, it might be helpful to look into parameters, which reflect more complex characterizations, as e.g. the amplitude variation or the relation of the power in different frequency bands (MQ_REL_). Since these parameters were newly developed in investigations with Parkinson patients without tremor [12], those parameters were not evaluated before in this context.

In the present study, the oscillatory parameters as the amplitude variation and the power-frequency ratio MQ_REL_ showed differences for the MMGobl, not for the MMGtri and MTGtri.

Theoretical considerations to explain these results might be found in neuroscientific research. Therein, it is assumed that the more adaptation is required during a motor task, the more relevant the feedback-control gets [20,21]. During HIMA, the participant has to react to the force of the partner, which performs PIMA. The pushing partner just has to apply the force level onto the counterpart and has not to react kinesthetically as intense as during HIMA. Therefore, it might be conceivable that HIMA entails higher requirements regarding the adaptive processes based on kinesthetic perception compared to PIMA. Hence, the thereby assumed more difficult neural control might explain the briefer endurance time during the holding task.

External variations, as a varying external force impact, require a constant update of the sensorimotor system [22]. To execute motor tasks appropriately an adequate adaptability is necessary. With a more complex feedback control, the variability will be even higher as a sign of an adequate adaptation [23]. In the present study, the amplitude variation of the mechanical muscular oscillations of the abdominal external oblique muscle was to be found significantly higher during HIMA with 83 ± 15% variation during the 15s trials and 79 ± 16% during the fatiguing trials compared to PIMA with 73 ± 14% and 70 ± 12%, respectively (*p* = 0.013, *r* = 0.55; *p* = 0.007, *r* = 0.57). This might be a sign for higher adaptation processes and, therefore, for more complex control strategies during the HIMA.

Furthermore, the power-frequency-ratio of the power in the low frequency range of 3 to 7 Hz related to the power in the area of 3 to 12 Hz might reflect more complex control strategies if the lower frequencies are represented more pronounced. In the present study, the power in the frequency range of 3 to 7 Hz amounted to 25 ± 15% during HIMA in the fatiguing trials and 17 ± 10% during PIMA (*p* = 0.002, *r* = 0.64) for the MMGobl. This might point out that the power distribution is changed during HIMA in a broader range, which possibly could enable the subject to react more appropriate to external changes as they were applied by the partner, who initiated and controlled the force level.

These considerations, of course, are still speculative. However, due to the high effect size and reasonable neuroscientific explanations, they seem to be conceivable and not that unlikely.

In looking more precisely at the performance of tasks HIMA and PIMA, we assume that HIMA is closer to eccentric muscle action and PIMA reflects rather the concentric muscle action – both in the sense of a “stopped” eccentric or concentric muscle action, respectively. This was hypothesized by Garner et al. [2] However, they rejected the hypothesis since they found no significant differences between HIMA und PIMA concerning the parameters EMG amplitude and frequency. As mentioned above, probably other parameters of the muscular output, as e.g. the amplitude variation or the power frequency ratio, respectively, could be more suitable to reflect the neuronal control processes more precisely.

During HIMA, the participant just should react to the partner, but in case of exceeding the maximal holding endurance under submaximal intensities, this partner would yield in direction of eccentric muscle action, thus, the muscle would lengthen. During PIMA, however, the participant works in direction of concentric muscle action. In case of a declining resistance offered by the partner in the course of time, the pushing subject would shorten its muscle in order to maintain the reaction force level, thus, would perform concentric muscle action. This leads to the assumption that HIMA might reflect the processes of eccentric and PIMA those of concentric muscle action. This could further underpin the hypothesis of more complex control strategies during HIMA, since several investigations suggest a higher complexity of the neural control during eccentric muscle actions [24–27]. This is not at least based on the findings, that during eccentric muscle action the cortical potential is higher compared to the concentric contraction [28]. However, the muscular activity measured by EMG is higher during concentric muscle action compared to eccentric one [26,28–35]. The results that, while performing PIMA during the fatiguing trials, the normalized amplitude tends to be higher and the power in the frequency range of 8 to 15 Hz is significantly higher for the MMGobl (not for MMGtri and MTGtri) might, therefore, support the assumption, that PIMA is closer to concentric muscle action. Reversely, during HIMA, the power in the same frequency range is lower, which might reflect the lower muscular activity during an eccentric muscle action. If a higher supraspinal activity is in fact apparent during HIMA remains open. This question has to be investigated by using methods, which are able to capture the supraspinal processes. Measurements using EEG, EMG and MMG were performed in our Neuromechanics Lab, but the evaluation remains.

### Considerations concerning the different findings between MMGtri, MTGtri and MMGobl

It has to be questioned, why the signals of the triceps brachii and its tendon did not show any significant differences, although this muscle group of the upper arm had to execute the motor tasks with respect to the elbow joint. The most conceivable reason might be that the forearms of both partners were coupled. Thereby, a synchronized mutual rhythm arises between both partners [8,9]. A coupling like this can occur basically in different ways [13]: Firstly, a master slave relation might arise, whereby one partner is active and the other one is passive. Since both neuromuscular systems are active, this coupling manner can be ruled out here. Secondly, both partners could agree to a mutual rhythm, whereby a distinctive interaction frequency would be generated. Thirdly, a leader-follower relation might arise, whereby one partner follows the other one. Since in the present setting the pushing partner initiates the force and the holding partner should react to this, we assume that a leader-follower constellation arises. For further discussion and mathematical reasoning see Schaefer & Bittmann [8,9]. Thereby, the oscillations might have been transferred through the interface onto the other partner. Since one partner performed HIMA while the other one performed PIMA, the specific characteristics of each tasks might have been eliminated in the MMGtri and MTGtri signals. The external abdominal oblique muscle was the most distal recorded muscle from the coupling point and, therefore, not directly coupled between the partners. This might be further supported by a study in the same setting, whereby the oscillations of the MMGtri, MTGtri and MMGobl signals showed interpersonal coherent behavior [9]. The coherence was highest for MTGtri and MMGtri. MMGobl showed also significant coherence, however, the duration of coherent phases was briefer.

In the former study concerning the two forms of isometric muscle action during single measurements [1], the power in the frequency range of 8 to 15 Hz and 10 to 29 Hz was to be found higher during HIMA compared to PIMA for the MTG signal of triceps tendon. This indicates that also the executing muscle-tendon structures show differences with regard to HIMA and PIMA if the coupling is not present. However, in the study of Rudroff et al. [5] investigating both motor tasks during muscular action of the elbow flexors at 20, 30, 45 and 60% of the MVIC using EMG, the power in the range of 10 to 29 Hz towards the end of measurement behaved reversed with higher amounts in PIMA compared to HIMA. This is consistent with the here presented results that the power in the frequency range of 8 to 15 Hz was higher during PIMA vs. HIMA for the MMGobl. The contrary results that in the single study the MTGtri showed higher power during HIMA might be due to the setting or due to the regarded structure, respectively. It is conceivable that the tendon oscillations’ might behave differently since three muscle heads insert into the triceps tendon. This could result in a deviating oscillating behavior. Further investigations have to examine this assumption.

## Conclusion

In the present setting, two participants interacted isometrically, whereby one partner performed the holding isometric tasks and the other one performed the pushing isometric tasks. The results showed that thereby differences between HIMA and PIMA occur independent of the physical conditions of the subjects, especially concerning the endurance capability, but also regarding the amplitude variation, the power in the frequency range of 8 to 15 Hz and the specific power frequency ratio of the MMG signal of the external abdominal oblique muscle. The MMG and MTG of the triceps brachii muscle and tendon, which are closer to the coupling interface between the partners, did not show this behavior. Future investigations have to examine whether or not the results can be supported, especially concerning the newly applied analysis of amplitude variation and the specific power-frequency ratio.

Also with regard to other investigations on this topic so far, some of the results support the assumption of two different forms of isometric muscle action, a holding and a pushing one. We suggest that the holding form requires higher neuronal control strategies, since a more adaptive functionality is demanded thereby. Probably, those higher requirements for adaptive processes could be an explanation of the observed reduced submaximal isometric endurance in HIMA. This is close to the considerations and the concept of the Adaptive Force [36,37]. Thereby, the neuromuscular system has to adapt isometrically to an increasing external force. If the maximal holding force is reached, the subject merges into an adapting eccentric muscle action. There are hints, that the Adaptive Force is vulnerable for disturbances of the neural system, as e.g. mental or olfactory inputs (publication in work).

Therefore, the holding isometric muscle action might adopt a specific status in sports and movement science, but also in neuroscience and medicine. As a first step, the two forms of isometric muscle action should be approved by additional research. In case of scientific confirmation, a gap in motor science should be filled since they are not considered therein until now.

## List of abbreviations

ACC: Acceleration sensor
CV: Coefficient of Variation
f: Female
HIMA: Holding Isometric Muscle Action
m: Male
MVIC: Maximal voluntary isometric contraction
MMG: Mechanomyography
MMGtri: Mechanomyography of triceps brachii muscle
MMGobl: Mechanomyography of the abdominal external oblique muscle
MQ_REL_: Arithmetic mean of the power in the frequency range of 3 to 7 Hz related to the power in the range of 3 to 12 Hz.
MTG: Mechanotendography
MTGtri: Mechanotendography of the tendon of the triceps brachii muscle
PIMA: Pushing Isometric Muscle Action
SD: standard deviation

## Declarations

### Consent for publication

The persons seen in Fig. 1 gave they written consent for publication.

### Availability of data and materials

The datasets used and/or analyzed during the current study available from the corresponding author on reasonable request.

### Authors’ contributions

Both authors (LS and FB) contributed to the conception and design of the study. LS perfomed the acquisition and analysis of data. In the process of interpretation and discussion of data both authors were participated. LS drafts the manuscript and it was critically revised by FB. Both authors gave their final approval of the version to be published.

## Acknowledgements

Not applicable.

## Supplementary information

S1 Table. Values of normalized amplitude.

S2 Table. Values of mean frequency.

